# Adaptation of *Pseudomonas aeruginosa* to repeated invasion into a commensal competitor

**DOI:** 10.1101/2024.03.19.585690

**Authors:** Rachel M. Wheatley, Liam P. Shaw, Sarah Shah, Selina Lindon, R. Craig MacLean

## Abstract

The host-associated microbiome is an important barrier to bacterial pathogen colonization and can mediate protection through a variety of mechanisms. We wanted to investigate the potential consequences of selection imposed by commensal bacterial competitors on an invading bacterial pathogen. To do this, we tested the ability of the opportunistic pathogen *Pseudomonas aeruginosa* to invade pre-established communities of an abundant commensal bacterium in the human microbiome, *Staphylococcus epidermidis*. We passaged ten independent lines of *P. aeruginosa* through daily invasion into a pre-established *S. epidermidis* population (coculture evolved lines), alongside daily passage through monoculture conditions (monoculture evolved lines). The monoculture evolved lines showed strong parallel evolution in the Wsp (Wrinkly spreader phenotype) signal transducing system involved in biofilm formation, and significantly elevated biofilm formation. On the other hand, adaptation to *S. epidermidis* occurred via mutations in a diverse set of genes, and the coculture evolved lines showed much weaker evidence for parallel evolution, suggesting that the selective pressure imposed by competition with *S. epidermidis* is more complex than the pressure imposed by culture conditions. Interestingly, the elevated biofilm formation phenotype seen in the monoculture evolved lines was not observed in the lines evolved in the presence of *S. epidermidis*, raising the question of whether enhanced biofilm formation did not evolve with *S. epidermidis* present because it was not beneficial, or because *S. epidermidis* may be able to restrict this evolutionary path by inhibiting biofilm formation.

## Introduction

Bacterial pathogens invading a human host will be exposed to a variety of different selective pressures. These selective pressures can drive adaptation, and in turn, shape the genomes of pathogen populations (*1, 2*). One important barrier to bacterial pathogen colonization is the host-associated microbiome, which can mediate protection against pathogen colonization via a number of direct and indirect mechanisms (*2, 3*). These mechanisms include through competition for space and nutrients or the production of inhibitory molecules (*4*). Successful colonization requires pathogens to successfully invade these host-associated microbial communities.

To gain a deeper understanding of how bacterial pathogens may adapt to commensal competitors during microbiome invasion, we use experimental evolution to explore the potential consequences of selection imposed by a pre-established commensal bacterial population. We use the opportunistic pathogen *Pseudomonas aeruginosa* along with an abundant commensal bacterium in the human microbiome *Staphylococcus epidermidis* as the model system for this work. *P. aeruginosa* is a major causative pathogen of infections in niches where *S. epidermidis* is an abundant commensal colonizer, primarily the skin and respiratory system (*5, 6*).

Rapid adaptation to the host environment is likely especially important for environmental opportunistic pathogens like *P. aeruginosa*, for whom the human host environment represents one of many niches it has the potential to survive in (*7-10*). *P. aeruginosa* is frequently implicated in chronic wound infections (*11*), serious short-term respiratory infections (*12*), and can also colonise the lungs of patients with bronchiectasis or cystic fibrosis resulting in recurrent long-term infections (*13*). *S. epidermidis* is suggested to contribute to pathogen colonisation resistance in the microbiome through a number of mechanisms (*14, 15*), including via the secretion of bacteriocins with inhibitory activity (*16, 17*) and via proteases that can inhibit biofilm formation, as demonstrated with *Staphylococcus aureus* (*18*). The interaction between *P. aeruginosa* and *S. epidermidis* is an important system to understand because *S. epidermidis* is a key commensal colonizer in *P. aeruginosa* infection niches, and similarly to as with *S. aureus, S. epidermidis* may also be implicated in polymicrobial infection settings with *P. aeruginosa (19, 20*).

In this study, we used experimental evolution to investigate the response of *P. aeruginosa* to repeated invasion into pre-established populations of *S. epidermidis*. We passaged ten independent lines of *P. aeruginosa* through daily invasion into pre-established cultures of *S. epidermidis* (coculture evolved lines), alongside daily passage through monoculture conditions (monoculture evolved lines). This experimental design gives *P. aeruginosa* the opportunity to adapt to *S. epidermidis*, while limiting *S. epidermidis* evolution through dilution with a daily refreshed population. We combined genome sequencing with phenotypic assays of the evolved *P. aeruginosa* populations to characterize the response of *P. aeruginosa* to repeated invasion into this commensal competitor.

## Methods

### Invasion assay of *P. aeruginosa* into *S. epidermidis*

To characterize the ability of *P. aeruginosa* to invade a pre-established culture of *S. epidermidis*, we used a green fluorescent protein (GFP)-tagged strain of *P. aeruginosa* strain PAO1 (PAO1-GFP) (*21*) (Supplementary Table 1). *S. epidermidis* (ATCC 14990) and PAO1-GFP were grown from glycerol stock on LB (Lysogeny Broth) Miller agar plates (Sigma-Aldrich) overnight at 37 °C. The following day, single colonies of both PAO1-GFP and *S. epidermidis* were inoculated into tryptic soy broth for overnight growth at 37 °C with shaking at 225 rpm in biological triplicate. The inner wells of a black-sided 96-well plate were filled with either 200 μL *S. epidermidis* overnight culture or 200 μL sterile tryptic soy broth, and inoculated with PAO1-GFP serially diluted to ∼5 x 10^5^ CFU/mL. To assess the growth of PAO1-GFP in either coculture with *S. epidermidis* or monoculture, GFP reads (excitation: 485/20, emission: 516/20) were taken at 10-minute intervals in a BioTek Synergy 2 microplate reader set to moderate continuous shaking for 24 hours. Relative fluorescence units (RFU; arbitrary units) over time were plotted using the Growthcurver package in R (*22*).

### Evolution experiment

The evolution experiment was performed as illustrated in Figure 1, adopting a similar experiment design as in (*23*). The passage experiment was performed in 5 mL round bottom culture tubes with snap cap lids containing 1 mL of tryptic soy broth. To initiate the experiments, *P. aeruginosa* (PAO1) and *S. epidermidis* were grown from glycerol stock on LB Miller agar plates overnight at 37 °C (Supplementary Table 1). Ten independent cultures of *P. aeruginosa* and *S. epidermidis* were initiated from single colonies into 1 mL tryptic soy broth and incubated overnight at 37 °C with shaking at 225 rpm. The following day *P. aeruginosa* was serially diluted to an inoculum of ∼5 x 10^5^ CFU/mL in either sterile tryptic soy broth (monoculture evolved lines) or in the pre-grown *S. epidermidis* cultures (coculture evolved lines) (Figure 1). The ten starting inoculums of *P. aeruginosa* were saved as glycerol stocks at -80 °C to use as paired ancestral strain comparisons for the evolved lines.

**Figure 1.**
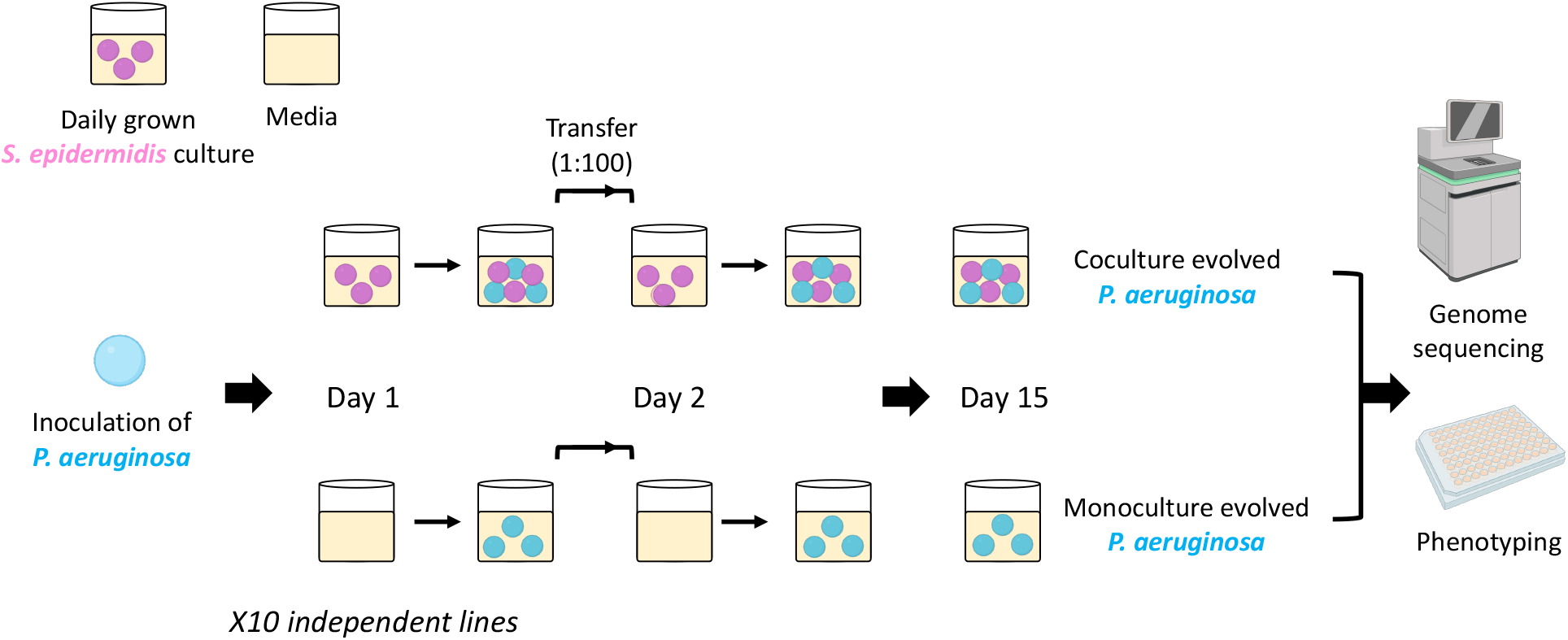
Evolution experiment design. *P. aeruginosa* was passaged in either monoculture or in coculture with *S. epidermidis* for 15 days of 24-hour incubations.

The evolution experiment cultures were incubated at 37 °C with shaking at 225 rpm for 24 hours, and fresh overnight cultures of *S. epidermidis* were prepared each day. For each passage, cultures were briefly vortexed and transferred at a 1:100 dilution into either the daily pre-grown *S. epidermidis* (coculture evolved lines) or sterile tryptic soy broth (monoculture evolved lines). This was repeated for 15 days. Samples were saved as glycerol stocks at the end of the 24h incubation for day 5, day 10, and the final day 15 passages. The samples from day 15 along with the saved starting *P. aeruginosa* inoculums were then grown from glycerol stock on LB Miller agar plates supplemented with 10 μg/mL vancomycin to select against *S. epidermidis*. Single *P. aeruginosa* colonies from each plate were used to produce a single isolate representative of each ancestral starting culture (x10), each coculture evolved line (x10), and each monoculture evolved line (x10).

### Phenotypic characterization of evolved isolates: growth assays

*P. aeruginosa* isolates were grown from glycerol stocks and single colonies inoculated into tryptic soy broth for 18-20h overnight growth at 37 °C with shaking at 225 rpm. Overnight cultures were diluted to an OD_595_ of ∼0.1 in either tryptic soy broth or in *S. epidermidis* spent media and placed within the inner 60 wells of a 96-well plate equipped with a lid. To assess growth, optical density (OD_595_) measurements were taken at 10-min intervals in a BioTek Synergy 2 microplate reader set to moderate continuous shaking for 22 hours, for two biological replicates per isolate. Growth in *S. epidermidis* spent media was used as a proxy for the coculture growth conditions. Spent media was prepared from overnight cultures of *S. epidermidis* via centrifugation at 16,000 g/min for 1 minute to pellet the cells followed by filter sterilisation of the supernatant through a 0.22 μM filter. Growth rate (mOD/min) was calculated as the maximum slope of OD versus time over an interval of ten consecutive readings. OD_595_ measurements over time were plotted using the Growthcurver package in R (*22*).

### Phenotypic characterization of evolved isolates: biofilm formation assays

Biofilm formation was quantified in microtiter plate assays as described in (*24*), with the alteration of tryptic soy broth to match the media used during experimental evolution. In brief, *P. aeruginosa* isolates were grown overnight from glycerol stocks and inoculated into tryptic soy broth in the inner 60 wells of a 96-well plate for overnight growth at 37 °C with shaking at 225 rpm, for six biological replicates per isolate. Overnight cultures were diluted 1:100 in fresh tryptic soy broth and incubated again overnight. For biofilm staining, plates were inverted and submerged in miliQ water twice, 125 μL 0.1% crystal violet solution was added to each well, the plate incubated for 15 minutes at room temperature, and then rinsed to remove excess dye before leaving to dry overnight. Biofilm formation was quantified by adding 125 μL of 30% acetic acid to each well for incubation at room temperature for 15 minutes, before transfer to a fresh plate and absorbance (OD_595_) quantified in the BioTek Synergy 2 microplate reader. Reads were background corrected against negative control wells and biofilm staining was compared between the starting culture isolates, monoculture evolved isolates, and coculture evolved isolates.

### Phenotypic characterization of evolved isolates: antibiotic resistance phenotyping

Antibiotic resistance of the *P. aeruginosa* isolates to ciprofloxacin, meropenem, and gentamicin was measured as minimum inhibitory concentration (MIC) assays via broth microdilution, as defined by EUCAST guidelines (*25*) with the alteration of the use of tryptic soy broth. Isolates were grown from glycerol stocks on LB Miller Agar plates overnight at 37 °C. Single colonies were then inoculated into tryptic soy broth for overnight growth at 37 °C with shaking at 225 rpm, after which overnight suspensions were serial diluted to ∼5 × 10^5^ CFU/mL. MIC was calculated using the 2-fold dilution series for meropenem (0.5 μg/mL - 8 μg/mL), ciprofloxacin (0.03125 μg/mL - 0.5 μg/mL), and gentamicin (0.5 μg/mL - 8 μg/mL). Growth inhibition was defined as OD_595_ < 0.200 and we calculated a single biologically independent MIC for each of the 30 *P. aeruginosa* isolates on each antibiotic. We analysed changes to antibiotic resistance for the pooled isolate groups of each culture condition (i.e. mean MIC of starting culture isolates, monoculture evolved isolates, and coculture evolved isolates).

### Genome sequencing

Bacterial gDNA was extracted from the 30 *P. aeruginosa* isolates (x10 starting cultures, x10 monoculture evolved, x10 coculture evolved) and gDNA concentration was quantified using the QuantiFluor® ONE dsDNA system kit (Promega), according to manufacturer’s instructions. The gDNA from 10 isolates for each culture condition was pooled in equimolar ratios and libraries were sequenced using the Illumina NovaSeq6000 platform using a 250 bp paired-end protocol, submitted for x510 depth sequencing. Library preparation and sequencing were performed by MicrobesNG (Birmingham, UK) according to their protocols.

### Variant calling

Read adaptor trimming was performed by MicrobesNG (Birmingham, UK) using Trimmomatic version 0.30 (*26*). The mean coverage of reads from *de novo* assembly confirmed a sequencing coverage of >500x (range: 669x-1090x). Variants were called using the breseq 0.38.1 pipeline in polymorphism mode (-p) aligned to a PAO1 reference genome (NCBI RefSeq accession NC_002516.2, accessed 23/06/23)(*27*), and annotations for mutations were added using snpEff v5.1d (*28*). Variants present in the monoculture or coculture pooled libraries were first identified by filtering the variants already identified in the pooled starting culture library. Then, variant positions exclusively present in either the monoculture or coculture pooled libraries were identified by filtering away positions also present in the alternative, to give a unique variant list for variant positions exclusively present in each endpoint. The pooled libraries were prepared by combining 10 isolates in equimolar ratios and we screened for variants present at a cut-off of at least 9% frequency in the pooled sequencing. Functional annotation was done using the Clusters of Orthologous Groups of proteins (COGs) database (*29*). Mutations on the *P. aeruginosa* PAO1 genome (NC_002516.2) were visualised using Proksee (*30*).

## Results

### Invasion of *P. aeruginosa* into a pre-established *S. epidermidis* population

We first set out to characterize the ability of *P. aeruginosa* to invade a pre-established *S. epidermidis* population by inoculating *P. aeruginosa* (PAO1-GFP) into an overnight culture of *S. epidermidis* and measuring growth over a 24-hour period. A pre-established *S. epidermidis* population could provide some resistance to *P. aeruginosa* invasion (Figure 2). The lag phase of *P. aeruginosa* increased from ∼8 hours to ∼14 hours in the presence of *S. epidermidis*, and growth of *P. aeruginosa* plateaued at ∼650 RFU compared to ∼1200 RFU that is reached in monoculture (Figure 2).

**Figure 2.**
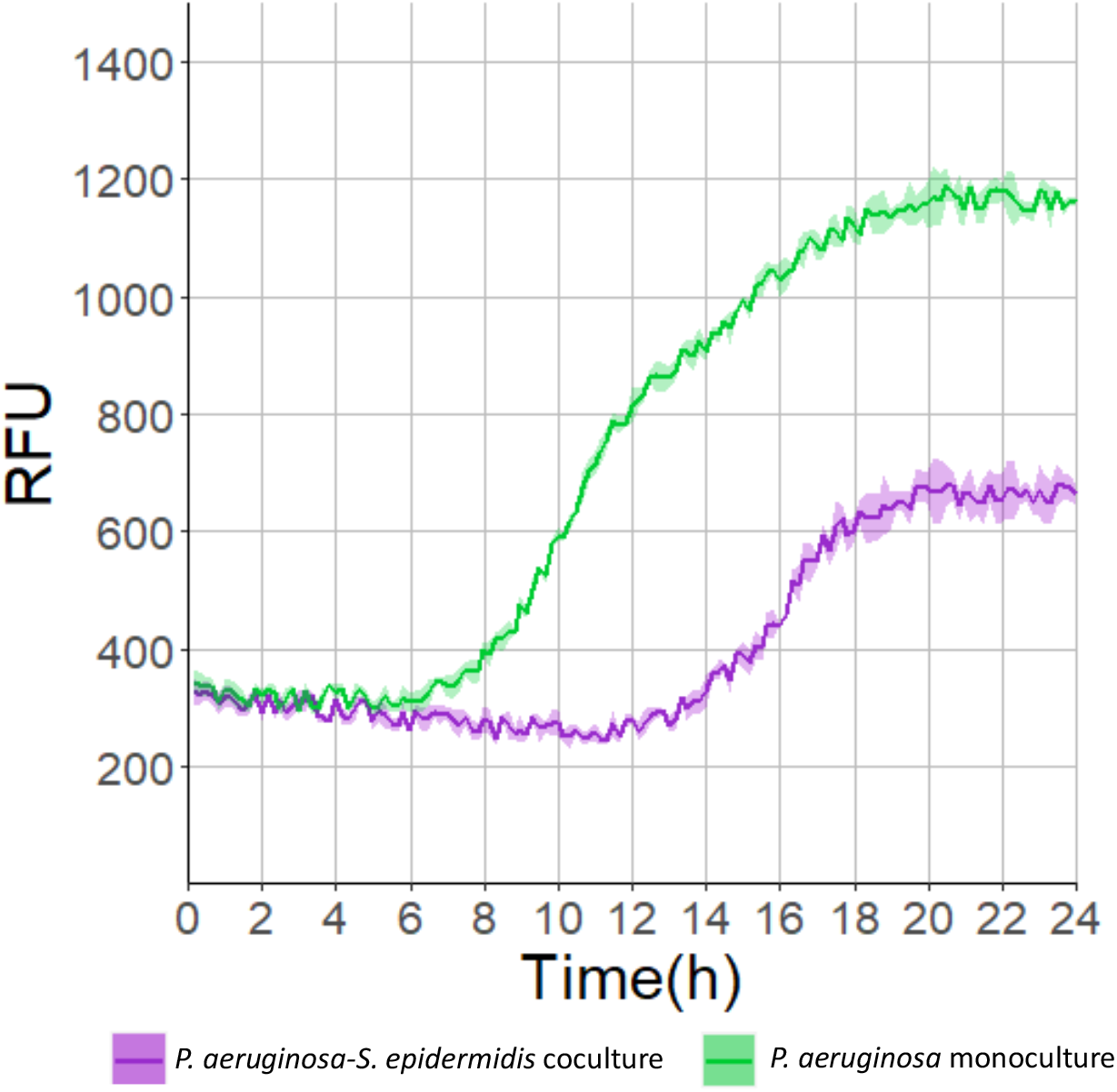
Characterisation of *P. aeruginosa* invasion ability into *S. epidermidis*. Growth of GFP-tagged *P. aeruginosa* (PAO1-GFP) when inoculated into either a pre-established population of *S. epidermidis* (coculture; purple) or in sterile tryptic soy broth (monoculture; green) as measured by relative fluorescence units (RFU). Graph shows the mean +/- s.d. calculated from three independent replicates.

### Phenotypic adaptation of evolved *P. aeruginosa* isolates

We characterized the growth, biofilm formation and antibiotic resistance phenotypes of the evolved *P. aeruginosa* isolates from the evolution experiment (illustrated in Figure 1) compared to the ancestral starting culture isolates. For the growth assays, *S. epidermidis* spent media was used as a proxy for the coculture conditions in which growth of *P. aeruginosa* could be measured via OD_595_. The coculture evolved isolates on average had a higher growth rate in *S. epidermidis* spent media compared to the ancestral starting culture isolates (Figure 3A, Figure 3B), but neither lag phase nor maximum OD_595_ were improved (Figure 3A). Growth in tryptic soy broth remained very similar for both the coculture evolved isolates and the ancestral starting culture isolates (Figure 3C), suggesting there was minimal adaptation to the media itself during the evolution experiment.

**Figure 3.**
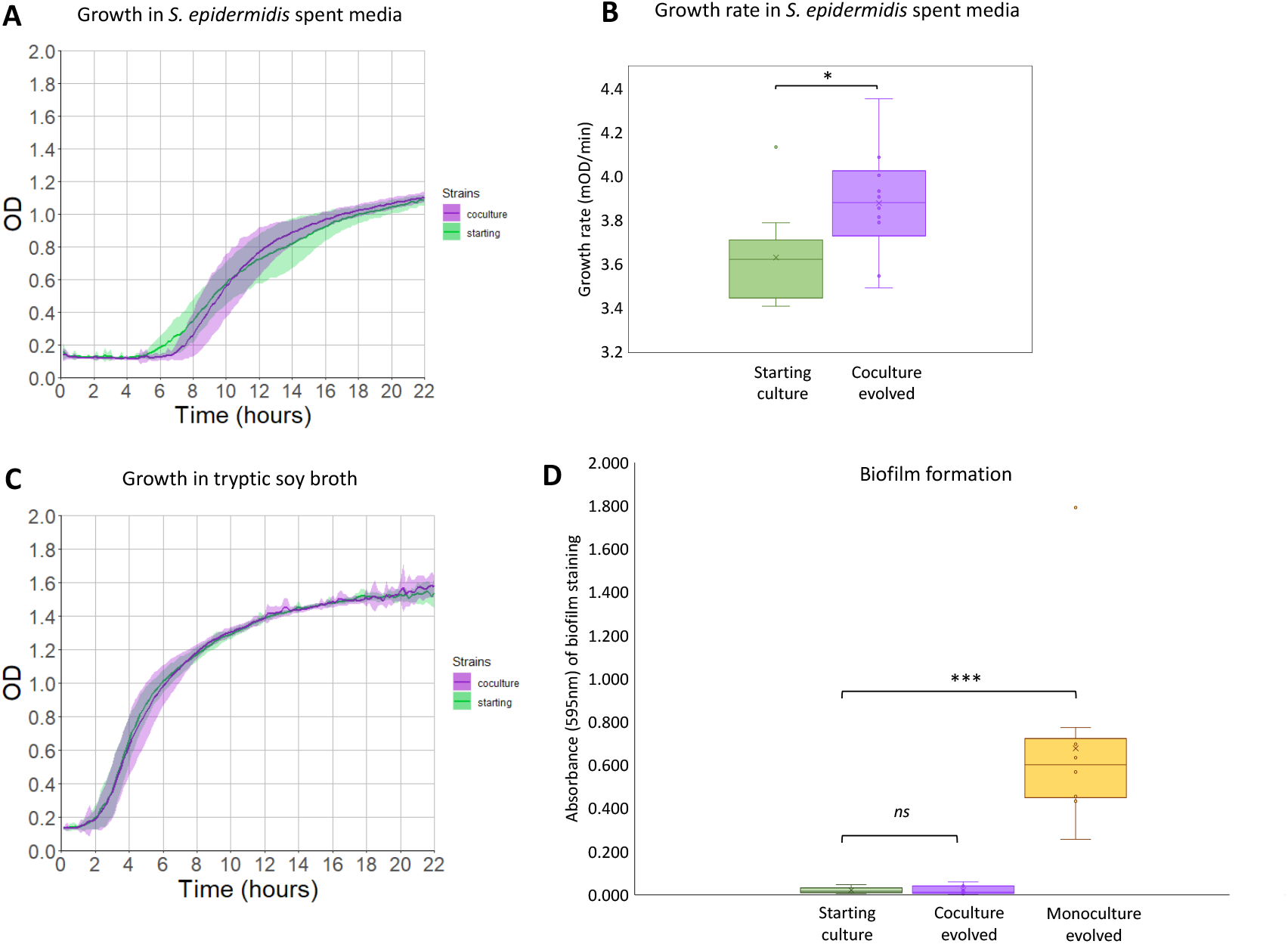
Phenotypic characterisation of evolved *P. aeruginosa* isolates. A) Growth of *P. aeruginosa* isolates from starting cultures (green) and coculture evolved endpoints (purple) in *S. epidermidis* spent media. Plot shows growth curve data as measured by OD_595nm_ from two replicate measurements of each isolate (apart from one of the coculture evolved isolates for which only one measurement was obtained) plotted as the mean +/- s.d. B) Growth rate (Vmax) of *P. aeruginosa* isolates from starting cultures (green) and coculture evolved endpoints (purple) in *S. epidermidis* spent media. C) Growth of *P. aeruginosa* isolates from starting cultures (green) and coculture evolved endpoints (purple) in tryptic soy broth as measured by OD_595nm_. Plot shows growth curve data as measured by OD_595nm_ from two replicate measurements of each isolate plotted as the mean +/- s.d. D) Absorbance (595nm) of biofilm staining obtained in the biofilm formation assay with the starting culture (green), coculture evolved (purple), and monoculture evolved (yellow) *P. aeruginosa* isolates. The means for each isolate as calculated from six biological replicates as plotted on the box plots. ns; non significant. *; significant at p < 0.05. ***; significant at p < 0.001.

Significant biofilm formation was noticeable when growing the monoculture evolved isolates, making it difficult to compare growth with our growth assay protocols. Biofilm formation is a phenotype of significant clinical interest because biofilms can act as a diffusion barrier, and contribute to a large proportion of recalcitrant hospital infections (*31, 32*). Using a biofilm formation assay we were able to validate that the monoculture evolved isolates had significantly elevated biofilm formation compared to the starting culture isolates (unpaired one-tailed t-test, p < 0.001), but this enhanced biofilm formation ability was not observed in the coculture evolved isolates (Figure 3D).

While *S. epidermidis* is not known to produce any antibiotics, one way *S. epidermidis* has previously been identified to contribute resistance to pathogen colonization is through the secretion of molecules (e.g. bacteriocins) with inhibitory activity (*16, 17*). General defense mechanisms of *P. aeruginosa* include efflux pump expression, and efflux pumps can transport many substrates including secreted molecules, metabolites, and antibiotics (*33, 34*). To this end, we were interested in whether the consequences of selection imposed by *S. epidermidis* included any changes in antibiotic susceptibility in *P. aeruginosa*. We measured the minimum inhibitory concentration (MIC) of each *P. aeruginosa* isolate to three different antibiotics (ciprofloxacin, meropenem, and gentamicin) representing three different classes of antibiotics. The MICs of both the coculture evolved isolates and monoculture evolved isolates did not differ significantly to the starting culture isolates (Supplementary Figure 1).

### Genome sequencing of evolved *P. aeruginosa* isolates

To understand the genomic basis of adaptation, we analysed the variants present in the coculture and monoculture evolved isolates (Figure 4) (Supplementary Table 2). In the pooled coculture evolved isolates, a total of 12 variants were identified (Supplementary Table 2), of which 6 variants were non-synonymous mutations. In the pooled monoculture evolved isolates, a total of 14 variants were identified (Supplementary Table 2), of which 7 variants were non-synonymous mutations.

**Figure 4.**
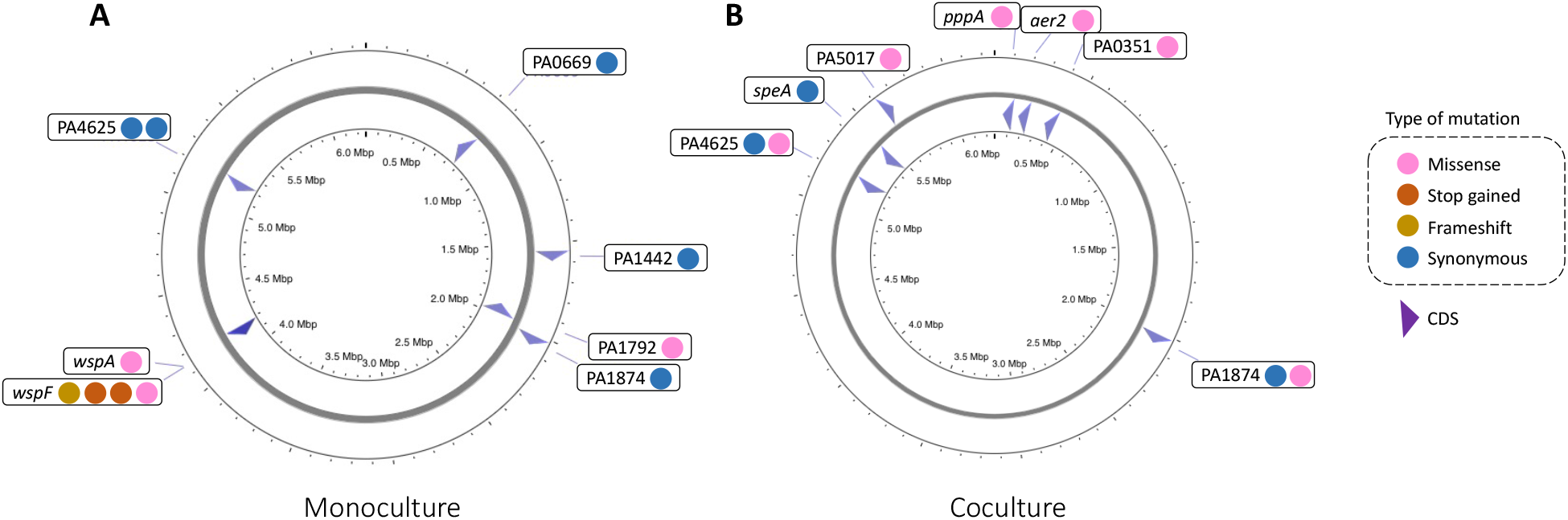
Map of mutations. A map of the protein changing (e.g. missense, frameshift, or gain of stop codon) and synonymous mutations identified in the pooled sequencing of the monoculture or coculture evolved isolates. This figure was created using a map of the *P. aeruginosa* PAO1 genome (NC_002516.2) in Proksee (*30*). Intergenic variants are excluded from this map.

In the coculture evolved isolates, non-synonymous mutations were identified in PA0176 (*aer2*), PA0351, PA1874, PA5017 (*dipA*), PA4625 (*cdrA*) and PA0075 (*pppA*) (Figure 4). PA0075 (*pppA*) is predicted to encode a regulatory component of the type VI secretion system (T6SS) in *P. aeruginosa* (*35*). PA0176 (aer2) is predicted to encode a methyl-accepting chemotaxis protein (Aer2), which senses oxygen and mediates stress responses, virulence and chemotaxis behaviour (*36*). PA5017 (*dipA*) is predicted to encode the phosphodiesterase DipA that contributes to biofilm dispersion by modulating c-di-GMP and polysaccharide levels (*37*).

In the monoculture evolved isolates, four non-synonymous mutations were identified in PA3703 (*wspF*) and one mutation was identified in PA3708 (*wspA*). The other non-synonymous mutations were in PA4625 (*cdrA*) and PA1792. Both PA3703 (*wspF*) and PA3708 (*wspA*) encode components of the Wsp (Wrinkly spreader phenotype) signal transducing system, a two-component system that is involved in biofilm formation (*38-40*). PA4625 (*cdrA*), in which non-synonymous mutations were identified in both the monoculture evolved and coculture evolved isolates, encodes a type V protein secretion system complex. Presence in both the monoculture and coculture evolved lines presumably indicates a role in adaptation to common laboratory conditions. Similarly, synonymous mutations were found in PA1874 in both the monoculture evolved and coculture evolved isolates.

## Discussion

In this work we used experimental evolution to investigate the adaptation of *P. aeruginosa* to repeated invasion into a commensal competitor, *S. epidermidis. S. epidermidis* is an abundant commensal in two significant infection niches for *P. aeruginosa*, the skin and the respiratory tract. Before starting our experimental evolution, we confirmed that a pre-established *S. epidermidis* population was able to provide some resistance to invasion by *P. aeruginosa* (Figure 2). Then, we passaged ten independent lines of *P. aeruginosa* through daily invasion into a pre-established *S. epidermidis* population, alongside daily passage through monoculture conditions. Selection in monoculture led to strong parallel evolution in the Wsp system, a two-component system that is involved in biofilm formation (Figure 4A). Phenotypic characterization of these monoculture evolved isolates confirmed significantly elevated biofilm formation (Figure 3B). Our phenotypic characterization of the coculture evolved isolates suggested that while *P. aeruginosa* was able to overcome some of the growth inhibition imposed by *S. epidermidis*, as indicated by an increased growth rate in *S. epidermidis* spent media (Figure 3B), adaptation to *S. epidermidis* occurred by mutations in a diverse set of genes (Figure 4B). The coculture evolved lines showed much weaker evidence for parallel evolution, suggesting that the selective pressure imposed by competition with *S. epidermidis* is more complex.

We identified a total of six non-synonymous mutations in the *P. aeruginosa* coculture evolved isolates (Figure 4B) (Supplementary Table 2). Three of these mutations were identified in genes annotated to function within signal transduction mechanisms (Table 1). Interestingly, one of these was in PA0075 (*pppA*), which is predicted to encode the regulatory component (PppA) of the type VI secretion system (T6SS) (*35*). The T6SS is used in many gram-negative bacteria to kill competing bacterial cells in densely occupied niches and functions by the injection of toxic effector molecules into other bacterial cells in a contact-dependant manner (*41-44*). Mutational inactivation of *pppA* has been shown to lead to increased T6SS activity (*45*). This suggests that the coculture evolved lines might fare better in competition with live *S. epidermidis* cells, rather than in this spent media proxy assay. We were additionally interested in whether passage through *S. epidermidis* cultures could select for any changes to biofilm formation capability or antibiotic susceptibility in *P. aeruginosa*, as these are two clinically relevant phenotypes for *P. aeruginosa* infections. We observed no significant difference in biofilm formation or in resistance to the antibiotics ciprofloxacin, meropenem and gentamicin between the starting cultures and the coculture evolved lines.

Looking at what is present in the coculture evolved lines is one way to understand the consequences of interaction with *S. epidermidis* but looking at what is absent from these lines is another way this can be achieved too. Selection in monocultures led to parallel evolution in the Wsp system, a two component system that is involved in biofilm formation (*38-40*), with five out of seven non-synonymous mutations being identified in either *wspF* or *wspA*. The monoculture evolved isolates demonstrated massively elevated biofilm formation compared to both the starting cultures and coculture evolved lines. That *wsp* mutations and enhanced biofilm formation were observed in the monoculture evolved isolates and not in the coculture evolved isolates raises the question of - *why not?* Is it that enhanced biofilm formation did not evolve with *S. epidermidis* present because it was not beneficial, or because *S. epidermidis* effectively blocks this evolutionary path by inhibiting biofilm formation? Although it is difficult to distinguish between these different possibilities, *S. epidermidis* is known to secrete a protease that can inhibit the biofilm formation of *S. aureus* (*18*), and it is possible a similar pathway may exist with *P. aeruginosa*.

## Supplementary Information

Supplementary Table 1. List of strains used in study.

Supplementary Table 2. Variants present in the monoculture and coculture evolved isolates.

Supplementary Figure 1. Minimum inhibitory concentrations of ciprofloxacin, meropenem and gentamicin.

## Acknowledgments

R.M.W and S.S. were supported by the Calleva Research Centre for Evolution and Human Sciences at Magdalen College, Oxford. R.M.W was supported by the George Grosvenor Freeman Fellowship by Examination in Sciences, Magdalen College (Oxford). L.P.S. was supported by a Sir Henry Wellcome Postdoctoral Fellowship (220422/Z/20/Z). This publication arises from research funded by the John Fell Oxford University Press Research Fund. We would like to thank Josh Thomas and Ashleigh Griffin (University of Oxford) for strains provided and used in this study. We would like to thank MicrobesNG (Birmingham, UK) for the generation and initial processing of sequencing data. Figures within this publication were created using Biorender.com (Figure 1).

## Author contributions

R.M.W. was responsible for conceptualization and funding acquisition. All authors contributed to investigation, methodology, and formal analysis. R.M.W., S.L. and R.C.M. contributed to data visualization. R.M.W., L.P.S. and R.C.M. contributed to writing and editing of the manuscript.

